# Proof-of-principle of NF1 Gene Therapy in plexiform neurofibroma mice models

**DOI:** 10.1101/2025.01.21.634081

**Authors:** Dhanushka Hewa Bostanthirige, Camille Plante, Jean-Paul Sabo Vatasescu, Maude Lévesque, Colin Poirier, Mathieu Deschenes, Benoit Chabot, Benoit Laurent, Sameh Geha, Jean-Philippe Brosseau

**Author notes:** Author for correspondence: Jean-Philippe Brosseau, Ph.D., Associate Professor, Department of Biochemistry and Functional Genomic, Centre de recherche du Centre Hospitalier Universitaire de Sherbrooke, Institut de recherche sur le Cancer de l’Université de Sherbrooke, Université de Sherbrooke. equal contribution (Jean-Paul Sabo Vatasescu, Maude Lévesque, Colin Poirier, Mathieu Deschenes).

## Abstract

Neurofibromatosis type I is a rare neurocutaneous syndrome characterized by the development of disfiguring neurofibroma tumors with unmet clinical needs. As Neurofibromatosis Type I is a monogenic disease, the development of gene therapy is highly attractive, but it is currently unknown if rescuing the *NF1* gene in established neurofibroma is sufficient for tumor regression. Here, we test this hypothesis by building two novel NF1 mouse models with reversible *NF1* expression. In the first model, the human *NF1* ^−/-^ Schwann cells named ipNF95.11b were genetically modified with a doxycycline-inducible full-length mouse *Nf1* gene. One month after cells implantation in the sciatic nerve, mice were split into 2 groups. Strikingly, all sciatic nerves from mice allowed to drink doxycycline water for one month display complete normalization of the sciatic nerve histologically (n=6 sciatic nerves) whereas 83% (5 out of 6 sciatic nerves) develop or maintain a neurofibroma when drinking regular water. In the second model, the human *NF1* ^+/-^ Schwann cells named ipNF95.11c were genetically modified with a doxycycline-inducible potent shRNA against the *NF1* mRNA transcript. Strikingly, doxycycline withdrawal after neurofibroma establishment allowed complete normalization (n=4 sciatic nerves), whereas all sciatic nerves showed evidence of neurofibroma when kept on doxycycline (n=4 sciatic nerves). Thus, we proof-of-principle NF1 Gene Therapy in plexiform neurofibroma mice models.

## INTRODUCTION

Neurofibromas are tumors arising in the peripheral nervous system that occur in nearly all patients with Neurofibromatosis Type I (NF1), a condition caused by mutations in the NF1 tumor suppressor gene within Schwann cells (1,2). Neurofibromas are disfiguring tumors that can severely affect daily life. Surgical options are limited, especially when numerous tumors are present or when they affect vital organs. Genetic analysis of Schwann cells from neurofibromas has shown that *NF1* is the sole gene consistently mutated across multiple samples (3), making NF1 a prime candidate for gene therapy. Gene therapy holds promise for NF1 patients, but its efficacy may take extensive research and significant funding to establish, potentially taking a decade to reach conclusions from phase 2 clinical trials (4,5). Despite NF1 mutations being implicated in neurofibroma formation, it remains untested whether restoring *NF1* function could disrupt tumor maintenance and serve as a definitive cure. This gap in knowledge is partly due to current NF1 mouse models, which involve permanent NF1 inactivation. Current models include mice with tissue-specific Nf1 ablation using Cre recombinase under a Schwann cell promoter (e.g. Krox-20, Dhh, Plp1, Hoxb7, Postn, So×10), leading to para-spinal neurofibroma with high penetrance within 4-12 months (6-11). Alternatively, graft models use human or mouse cells with biallelic *Nf1* or *NF1* inactivation in the vicinity of a sciatic nerve of immunocompromised mice. These models confirm neurofibroma development through histological analysis after 1-4 months (11-14). However, these models are unsuitable for testing the conceptual feasibility of NF1 gene therapy due to permanent gene inactivation. To address this, novel NF1 mouse models were developed with reversible NF1 expression to investigate whether the loss of NF1 is essential for neurofibroma maintenance.

## RESULTS

We began by developing a xenograft recapitulating plexiform neurofibroma (pNF) using human *NF1* ^−/-^ Schwann cells. Briefly, triple-immunocompromised NCG mice were implanted ipNF95.11b cells in the vicinity of injury-induced sciatic nerves. After three months, sciatic nerves were harvested and submitted to histological evaluation following guidelines from the literature (15,16). Compared to normal nerves, we consistently observed higher cellularity, disorganized neural architecture characteristic of neurofibroma and presence of thick collagen bundles (Fig. 1A, S1A). These neurofibroma characteristics were not consistently observed when performing a sham surgery or xenograft with normal or *NF1* ^+/-^ Schwann cells (Fig. 1A, S1B-D). To ensure that the developed pNFs originate from the implanted human Schwann cells, we performed immunohistochemistry using Schwann cell-specific (S100) and human-specific (Ku80) markers (Fig. 1A, S1A-D). Overall, pNF were only and consistently observed when ipNF95.11b cells were implanted (n=4 sciatic nerves per condition) (Fig. 1B). Of note, there was a similar high penetrance of pNF even at 1 month in NCG mice (5 out of 5 sciatic nerves) and nude mice (5 out of 6 sciatic nerves) (Fig. 1C, S1E-F). Thus, ipNF95.11b graft into nude mice represents a rapid and robust pNF assay.

**Figure 1.**
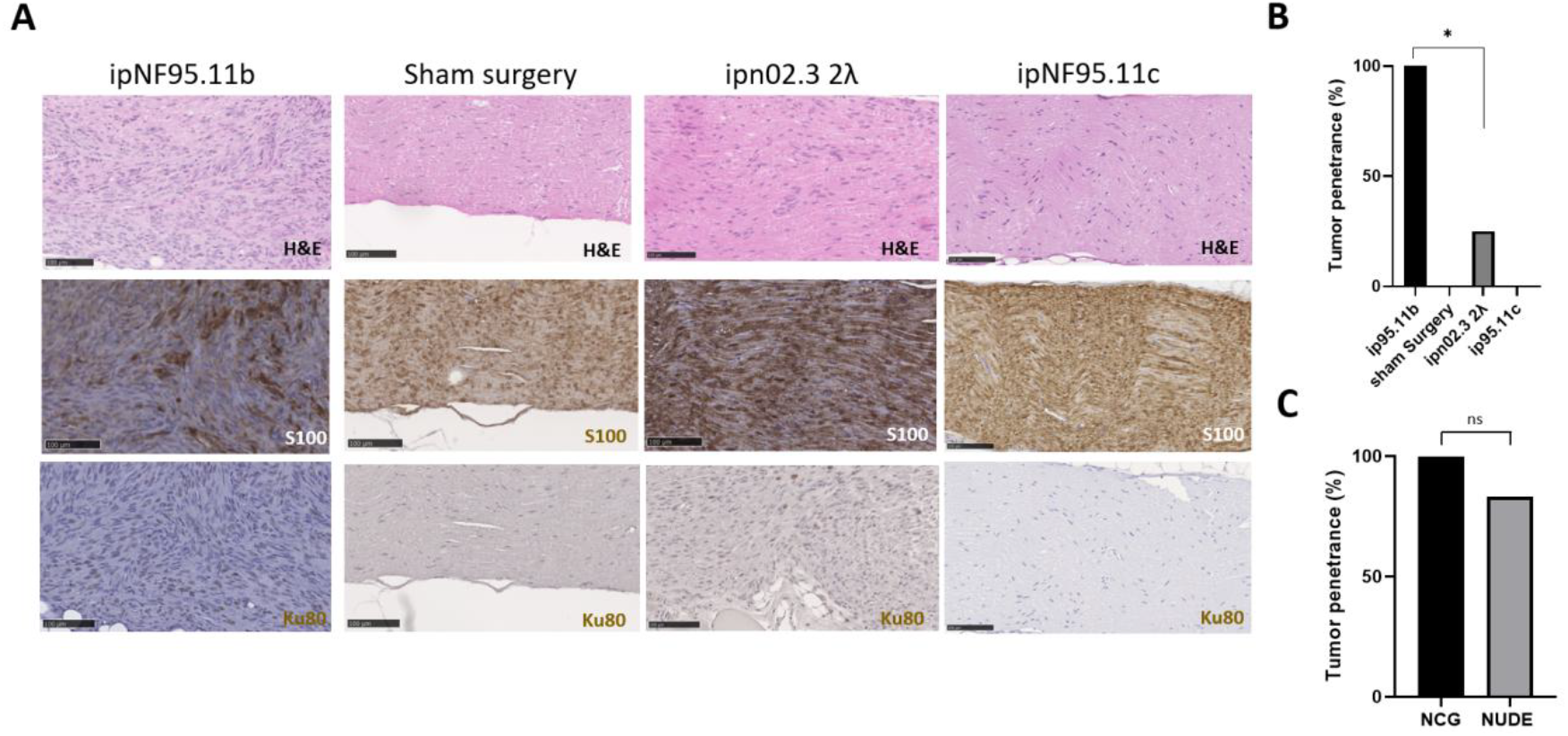
Development of the ipNF95.11b xenograft assay. A) Representative H&E (upper), S100 [(Schwann cell marker) (middle)] and Ku80 [(human-specific marker) (bottom)] and immunohistologies of sciatic nerve from NCG mice implanted with ipNF95.11b cells (first column), no cells (2^nd^ column) ipn02.3 2λ (3rd column) or ipNF95.11c (4^th^ column) and harvested after 3 months. The scale bar equals 100 μm. B) Tumor penetrance of the experiment in A (n=4 sciatic nerves per condition). C) Bar graph representing the comparison of tumor penetrance of the ipNF95.11b xenograft assay in NCG (n=5 sciatic nerves) and nude mice (n=6 sciatic nerves) at 1 month. Fischer exact t-test was used to measure statistical significance (* means p ≤ 0.05). ns means non-significant.

Next, we assessed whether the re-expression of NF1 in neurofibroma tumor cells was sufficient to alter tumor maintenance. We genetically modified the ipNF95.11b cells with a lentiviral vector expressing the mouse *Nf1* gene under the control of the inducible Tet-ON system (pLVX-TetOne-Nf1). To ensure that the system was working as expected, we validated Nf1 expression increased upon doxycycline induction *in vitro* by qPCR (Fig. 2A). We next performed an experiment where immunocompromised mice were implanted with ipNF95.11b-TetOne-Nf1 cells. After one month (the time to develop neurofibroma), mice were allowed or not to drink doxycycline water to respectively induce or not Nf1 expression (Fig. 2B). After 2 months, all mice were euthanized, sciatic nerves were harvested and processed for histology. Strikingly, no nerves (0 out of 6 sciatic nerves) from mice with tumors expressing exogenous *Nf1* show tumors (group 2) while sciatic nerves (5 out of 6 nerves) from control mice (group 1) still exhibited neurofibroma (Fig. 2C-D, S2A-D). These results demonstrate that neoplastic nerve tissue can be normalized by re-expression of NF1 in a pNF xenograft model. Importantly, we tested the capacity of ipNF.95.11b-TetOne-Nf1 cells to form pNF within 1 month (Fig. 2B, group 3). 100% (4 out of 4 sciatic nerves) classified histologically as pNF (Fig. 2D, S2E). The fact that we obtained similar results using the ipNF95.11b (Fig. 1C, S1F) and ipNF95.11b-TetOne-Nf1 without doxycycline (Fig. 2D, S2E) confirms minimal leakage of the Tet-ON system. Thus, it demonstrates that neoplastic nerve tissue can be normalized by re-expression of NF1 in a pNF xenograft model.

**Figure 2.**
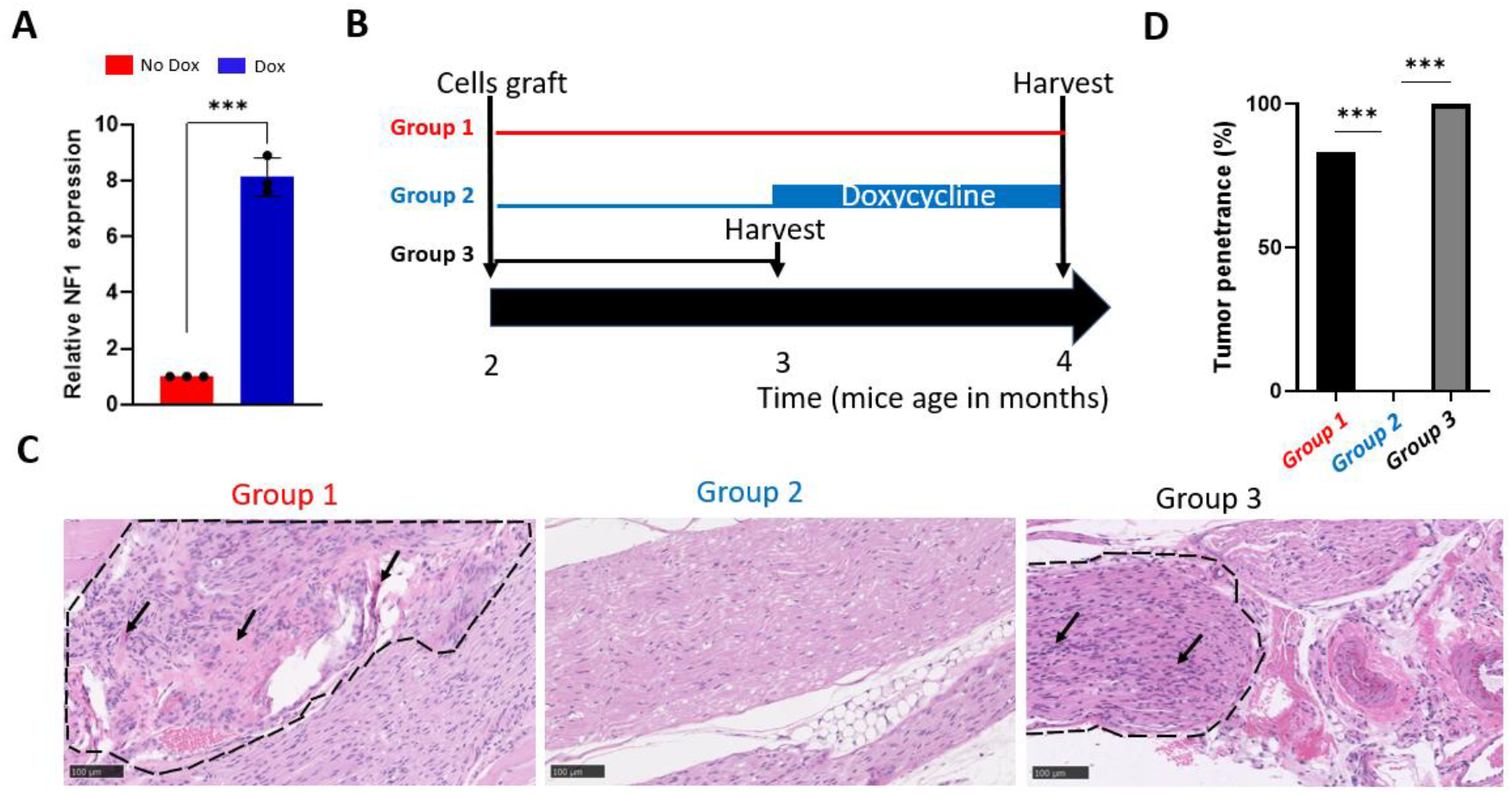
Re-expressing mouse Nf1 normalize neurofibroma in the ipNF95.11b_TetOne-Nf1 xenograft model. A) Validation of the Tet-ON inducible Nf1 expression system *in vitro* by qPCR. B) Schematic illustrating the experimental *in vivo* design to proof-of-principle NF1 gene therapy. C) Representative H&E for groups 1, 2 and, 3. Dash lines circumscribed the cellular and neurofibroma architecture area. Arrows point to collagen bundles. The scale bar equals 100 μm. D) Bar graph representing the comparison of tumor penetrance of group 1 (n=6 sciatic nerves), group 2 (n=4 sciatic nerves) and, group 3 (n=6 sciatic nerves). Fischer exact t-test was used to measure statistical significance (*** means p ≤ 0.01).

However, these encouraging results rely on the exogenous expression of mouse *Nf1*. To demonstrate the re-expression of human NF1, we developed a second xenograft strategy using the human *NF1* ^+/-^ Schwann cell line ipNF95.11c. We modified these cells to express a potent shRNA against NF1 (shNF1) under the inducible Tet-ON promoter. To ensure that the system was working as expected, we validated NF1 expression upon doxycycline induction *in vitro* by qPCR (Fig. 3A). We then performed an experiment where immunocompromised mice were implanted with ipNF95.11c_TetON-shNF1 and allowed to drink doxycycline for one month to knockdown NF1 expression and ultimately stimulate neurofibroma genesis. After one month, when tumors were fully established, mice were withdrawn water with doxycycline for 1 month to restore NF1 expression (Fig. 3B). Mice were euthanized, and sciatic nerves were harvested and processed for histology. We observed the tumor regression and tissue normalization compared to control animals, still under doxycycline water (Fig. 3C, S3A-B). Indeed, no nerves (0 out of 4 sciatic nerves) from mice with tumors expressing back NF1 show tumors (Fig. 3C-D, S3A) while all nerves (4 out of 4 sciatic nerves) from control mice still exhibited neurofibroma (Fig. 3C-D, S3B).

**Figure 3.**
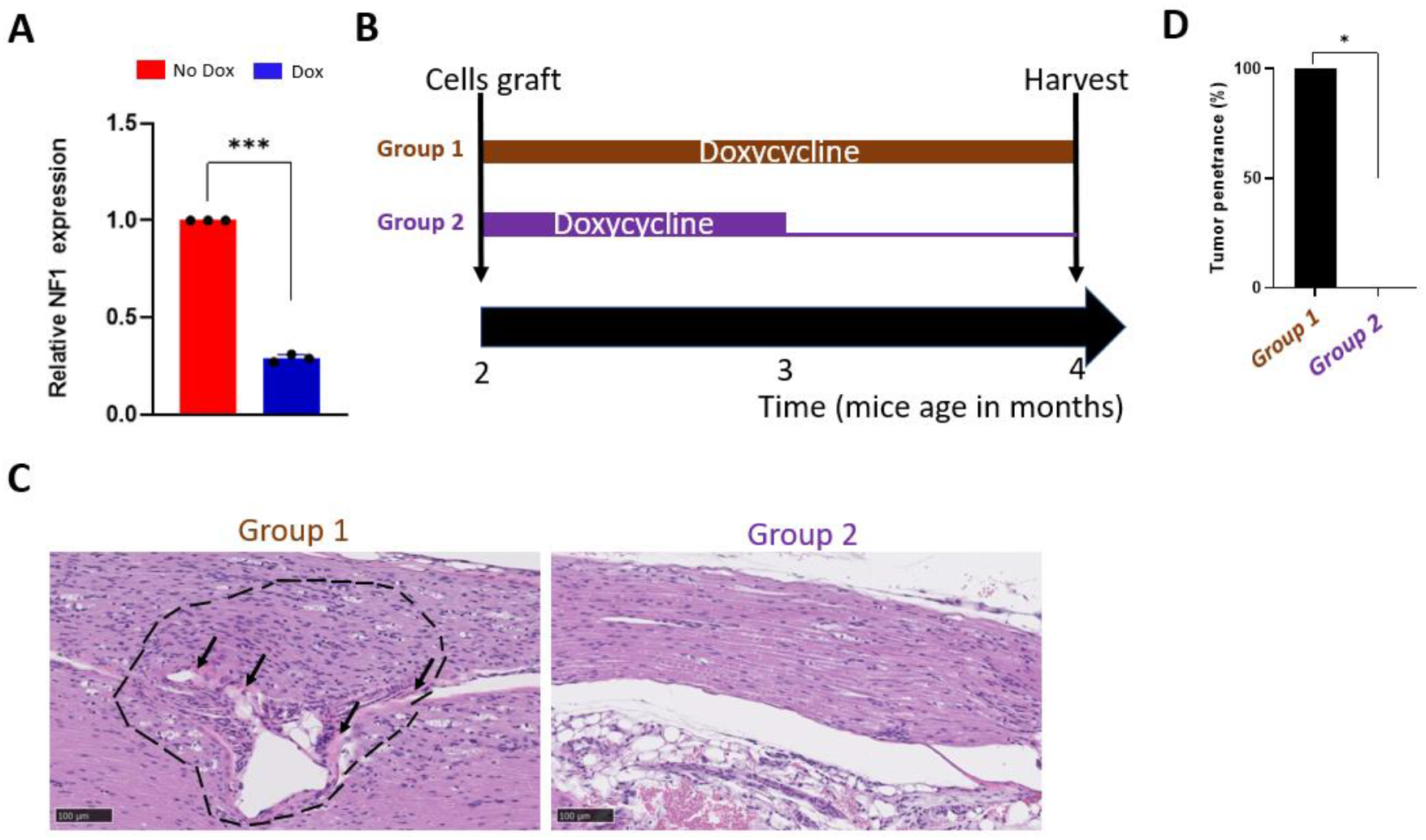
Re-expressing human NF1 normalize neurofibroma in the ipNF95.11c_TetON-shNF1_2 xenograft model. A) Validation of the Tet-ON inducible shNF1 expression system *in vitro* by qPCR. B) Schematic illustrating the experimental *in vivo* design to proof-of-principle NF1 gene therapy. C) Representative H&E for groups 1 and 2. Dash lines circumscribed the cellular and neurofibroma architecture area. Arrows point to collagen bundles. The scale bar equals 100 μm. D) Bar graph representing the comparison of tumor penetrance of group 1 and 2 mice (n=4 sciatic nerves per condition). Fischer exact t-test was used to measure statistical significance (* means p ≤ 0.05).

## DISCUSSION

There is significant enthusiasm in the NF1 scientific community for developing a gene therapy to reverse the clinical manifestation of NF1. However, proof-of-principle for NF1 gene therapy in animals for any NF1 clinical manifestation has yet to be demonstrated, partly due to the absence of suitable models. The workhorses of the field are tissue-specific *Nf1* knockout mouse models generating para-spinal neurofibroma with high penetrance (6-11), a rare human neurofibroma subtype. All these models rely on the same DNA alteration in *Nf1* exon 31 (17). Here, we took advantage of the commercially available ipNF95.11b to build novel mouse models developing pNF in the peripheral nerves as generally occurring in NF1 patients. Leveraging the Tet-On system, we have shown convincingly that re-expressing full-length human or mouse neurofibromin is sufficient to normalize peripheral nerves. Traditionally, neurofibromin was primarily viewed as a Ras GTPase-activating protein (GAP), and technical challenges in cloning the full-length 12kb NF1 gene into expression vectors led many researchers to use the NF1-GRD as a substitute. However, introducing NF1-GRD in *NF1* ^−/-^ Schwann cells only partly rescues the Ras-GTP levels (18) and cell proliferation (19), indicating that NF1-GRD may not be suitable for NF1 gene therapy. An alternative strategy was recently put forward. Around 20% of *NF1* mutations translate into pre-maturation stop codon mRNAs targeted for the nonsense mediated decay (NMD) mechanism, and hence lower NF1 expression. Ataluren is a small-molecule drug with proven efficacy in suppressing NMD. The group of Allan Jacobson hypothesized that ataluren treatment in tumor cells harboring an *NF1* mutation creating a premature stop codon would lead to an increase in NF1 expression and, ultimately, milder clinical manifestations in NF1 patients. Towards this goal, they demonstrated that mouse neuronal cells with an *Nf1* ^R683X/R683X^ mutation treated with ataluren can promote readthrough of the nonsense mutation at codon 683 of *Nf1* mRNA *in vitro* but only show a modest increase in NF1 expression level (20). *In vivo*, results using the Dhh-cre *Nf1* ^R683X/4F^ mice indicated so far that ataluren might slow the growth of neurofibromas and alleviate some paralysis phenotypes (21). Importantly, ataluren is not specific to NF1 mRNA and, hence, cannot be used to address the question of whether NF1 re-expression in established neurofibroma is sufficient to make tumor regress.

Another study investigated the concept of nonsense suppression in a novel NF1 animal model harboring a mutation creating a premature stop codon: the *NF1* ^R1947/+^ Ossabaw minipig model (22). In vitro, *NF1* ^−/-^ Schwann cells isolated from neurofibroma and treated with any established nonsense suppression drugs (ataluren, gentamicin, or G418) did not reliably increase NF1 expression although some reduction in MAPK signaling was observed. These findings highlight the challenges and potential strategies for developing effective gene therapies for NF1.

## CONCLUSIONS

We have established a proof-of-principle for NF1 gene therapy in plexiform neurofibroma mouse models. This breakthrough demonstrates the feasibility of extending NF1 gene therapy to address other clinical manifestations of the disorder. Ultimately, this research paves the way for developing pharmacological therapies that could benefit NF1 patients.

## EXPERIMENTAL PROCEDURES

### Mice husbandry

NCG and nude mice were purchased from Charles River. All mice were housed in the animal facility of the Institut de recherche sur le Cancer de l’Université de Sherbrooke, Quebec, Canada. All procedures have been approved by the Animal Research Ethics Committee of the Faculty of Medicine and Health Sciences (FMSS) of the Université de Sherbrooke and strictly follow the Canadian Council on Animal Care (CCAC) requirements. All mice were housed in hermetically sealed cages and placed on a support equipped with a ventilation system. All mouse strains were maintained in a room with 12/12 (day/night) light cycle with a temperature of 70-72°F

### Xenograft assay

Under constant isoflurane anesthesia, nude mice’s skin was cut open above the left quadriceps. The left sciatic nerve was located and damaged using a 27G needle. 1 × 10^6^ cells in 100 uL of L-15 culture media were implanted in the vicinity of the left sciatic nerve. The skin was sutured with monocryl 4-0 (absorbable). The procedure was repeated similarly for the right sciatic nerve. Finally, mice were allowed to recover on a warm pad until fully awakened and placed back into a cage.

### Cell lines

Ipn02.3 2λ, ipNF95.11b, ipNF95.11c were purchased from ATCC. Cells were passaged and expanded in DMEM + 10% FBS.

The pLVX-TetOne-Nf1 vector was constructed by cloning the Nf1 mouse open reading frame from the host vector (Genecopoeia; EX-Mm04084-M02) in place of the GFP open reading frame of the recipient pLVX-TetOne-GFP vector (addgene; #171123).

The pLKO-TetON-shNF1 was constructed by cloning the following DNA oligonuclotides (shNF1_FWD 5’- CCGGTAAGCGGCCTCACTACTATTTCTCGAGAAATAGTAGTGAGGCCGCTTATTTTTG-3’ and shNF1_REV 5’-AATTCAAAAATAAGCGGCCTCACTACTATTTCTCGAGAAATAGTAGTGAGGCCGCTTA-3’) into the recipient Tet-pLKO-puro vector (addgene; #21915).

Stable cell line establishment was performed by infecting recipient cells with lentiviral particles generated from the desired inducible vector and the following 3 packaging vectors : plp1 (addgene; #6097); plp2 (addgene; #6098); plp/VSVG (addgene; #6099) in HEK293T cells followed by puromycin selection.

### Doxycycline treatment

Doxycycline drinking water for mice was prepared by diluting 1g of doxycycline (Sigma, D9891) in 500 mL of autoclaved water containing 5% sucrose (Bioshop, SUC507). For in vitro studies, cells were treated with doxycycline at a final concentration of 2ug/uL and incubated for 3 days.

### Real-time PCR

RNA extraction was performed using the Trizol reagent following the manufacturer’s recommendation. The resulting total RNA extract was quantified using Nanodrop, and 150 ng from each sample were submitted to a reverse transcription reaction using Roche Transcriptor™ and N6 Roche random primers in a 10uL reaction. Finally, cDNA was diluted with 440 uL of RNase DNase free water and used as template for a qPCR reaction using NF1-specific primers (mouse-Nf1-FWD 5’-GCAACTTGCCACTCCCTACTGA, mouse-Nf1-REV 5’-ATGCTGTTCTGAGGGAAACGCT-3’; human-NF1-FWD 5’-AAAGGATCCCACTTCCGGTG-3’; human-NF1-REV 5’-CTTGGTCGCTCTCCCCACTA-

3’) and GAPDH as housekeeping gene (mouse-Gapdh-FWD 5’-GCGAGACCCCACTAACATCA-3’, mouse-Gapdh-REV 5’-GGCGGAGATGATGACCCTTT-3’) in the 2X SyBr Green mix buffer (Quantabio, 95054-02K) under the following cycling conditions: 95°C, 3 min; [(95°C, 15sec, 60°C, 30sec, 72°C, 30sec) X 50], 72°C, 30 sec. Relative expression was calculated using qBase (23).

### Histology

Mice were sacrificed under isoflurane, followed by CO^2^ asphyxia. Sciatic nerves were harvested and fixed in 10% formaldehyde (Sigma, HT501128-4L) for a minimum of 48h. Tissues were washed in 70% ethanol, paraffin-embedded, and cut at 5 um. H&E slides were prepared from the resulting formalin-fixed paraffin-embedded (FFPE) slides and scanned on Hamamatsu nanozoomer. Tissues were classified as pNF when the 3 histological criteria were observed: higher cellularity compared to normal nerves, disorganized neural architecture characteristic of neurofibroma and presence of thick collagen bundles.

Immunohistochemistry (IHC) of the sciatic nerves was performed based on the Vectastain ABC kit (Biolynx, VECTPK4000) and the DAB substrate kit peroxidase (Biolynx, VECTSK4103100) following the manufacturer’s recommendation. A boiling pH=6 acetate buffer as the antigen retrieval solution and the following primary antibodies [(anti-S100, GA50461-2); (anti-Ku80, NEB 2180S, 1:200)] and secondary antibody (anti-rabbit (Jackson Immunoresearch, 711-065-152, 1 :1000) were used. The pathologist (S.G.) was blinded to the mice experimental group when performing the histological review.

### Statistical analyses

Unless otherwise stated, a Fischer exact t-test was used to measure statistical significance (p ≤ 0.05).

## ACKNOWLEDGEMENT

We are grateful to the JPBrosseau lab members for proofreading. We thank Simon Roy from Civic Biosciences for his intellectual input on the design and production of the pLVX-TetOne-Nf1 vector. The Electron Microscopy and Histology research core of the Faculté de Médecine et des Sciences de la Santé at the Université de Sherbrooke for their histologies. We thank the RNomic platform for its PCR services. DHB is a recipient of the CRCHUS postdoctoral fellowship. JPB is a recipient of the New Investigator Award from the US Department of Defense and a FRQS J2 research scholar. This research was supported by operating grant # 942244 from the Cancer Research Society and grants from Association de la Neurofibromatose du Quebec since 2021.

## AUTHOR CONTRIBUTION

Conceptualization: JPB Experiments: DHB, CPlante

Preliminary results: JPSV, CPoirier, ML, MD, BL

Histological evaluation: DHB, JPB, SG Supervision: JPB, BL, BC

Draft writing: JPB Editing: JPB, BL, BC Funding: JPB

## DATA AVAILABILITY STATEMENT

Upon request at

