## Supplementary material for "Proof-of-principle of NF1 Gene Therapy in plexiform neurofibroma mice models": Suppl File 1

#### SUPPLEMENTARY FIGURE LEGENDS

**Figure S1. Histological characterization of the ipNF95.11b xenograft assay.** A-F) H&E ($1^{st}$ and $2^{nd}$ row), S100 [(Schwann cell marker) ($3^{rd}$ row)] and Ku80 [(human-specific marker) ($4^{th}$ row)] immunohistology of sciatic nerves from A-D) NCG mice implanted with A) ipNF95.11b (n=4 sciatic nerves), B) no cells (n=4 sciatic nerves), C) ipn02.3 2λ (n=4 sciatic nerves), D) ipNF95.11c (n=4 sciatic nerves), and harvested after 3 months and E) NCG (n=5 sciatic nerves) or F) nude mice (n=6 sciatic nerves), implanted with ipNF95.11b cells and harvested after 1 month. Scale bar equals 1 mm ($1^{st}$ row) or 100 µm ($2^{nd}$ – $4^{th}$ row).

**Figure S2. Histological characterization of the ipNF95.11b-TetOne-Nf1 xenograft assay.** A-E) H&E ($1^{st}$ and $2^{nd}$ row), S100 [(Schwann cell marker) ($3^{rd}$ row)] and Ku80 [(human-specific marker) ($4^{th}$ row)] immunohistology of sciatic nerves from nude mice implanted with ipNF95.11b_TetOne-Nf1 cells A-B) harvested after 2 months (n=6 sciatic nerves), C-D treated with doxycycline after 1 month and harvested after 2 months (n=6 sciatic nerves), E) harvested after 1 month (n=4 sciatic nerves). Scale bar equals 1 mm ($1^{st}$ row) or 100 µm ($2^{nd}$ – $4^{th}$ row).

**Figure S3. Histological characterization of the ipNF95.11c-TetON-shNF1 xenograft assay.** A-B) H&E ($1^{st}$ and $2^{nd}$ row), S100 [(Schwann cell marker) ($3^{rd}$ row)] and Ku80 [(human-specific marker) ($4^{th}$ row)] immunohistology of sciatic nerves from nude mice implanted with ipNF95.11c_TetON-shNF1 cells A) treated with doxycycline for 2 months and harvested at 2 months (n=4 sciatic nerves), B) treated with doxycycline for 1 month and harvested after 2 months (n=4 sciatic nerves). Scale bar equals 1 mm ($1^{st}$ row) or 100 µm ($2^{nd}$ – $4^{th}$ row).

### Figure S1

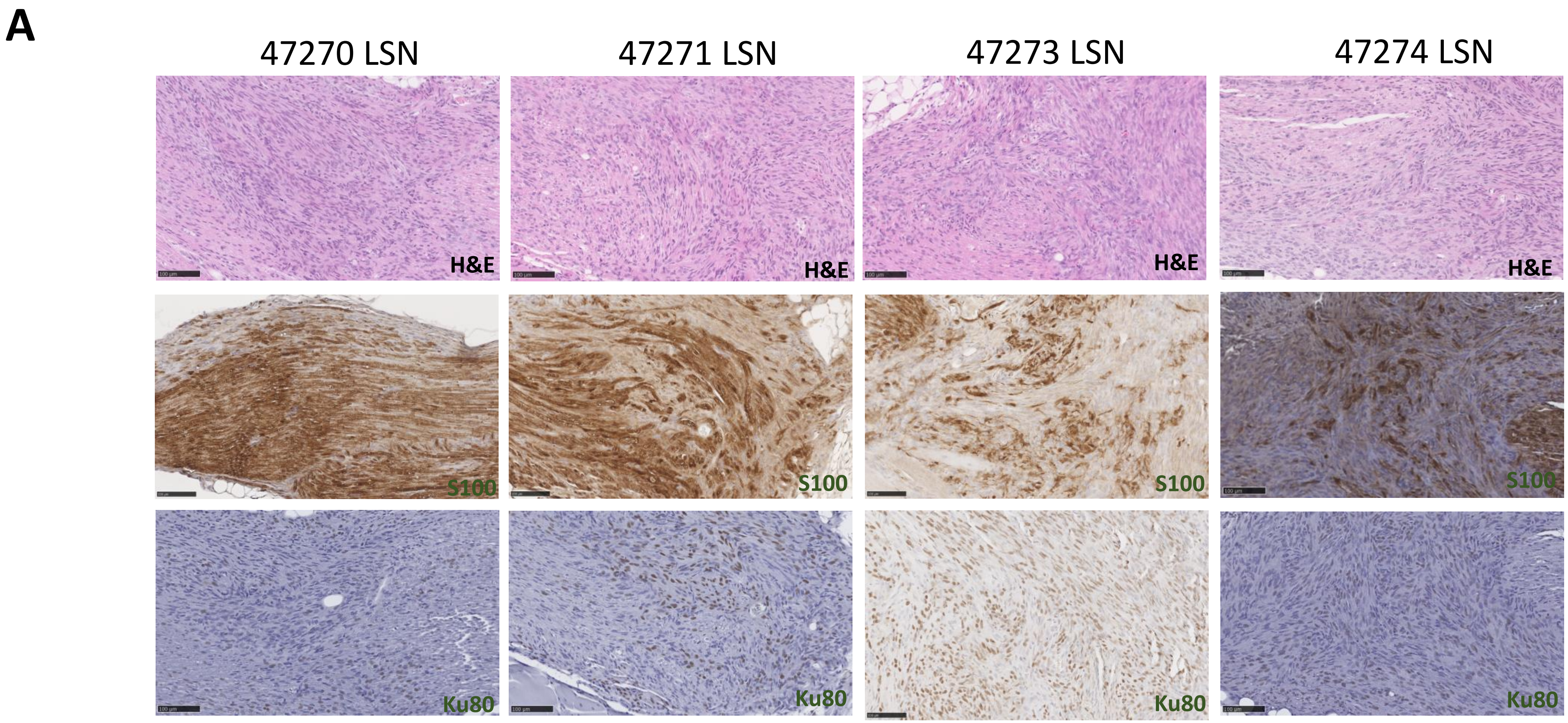
A
47270 LSN
47271 LSN
47273 LSN
47274 LSN
H&E
H&E
H&E
H&E
S100
S100
S100
S100
Ku80
Ku80
Ku80
Ku80

### Figure S1

## B

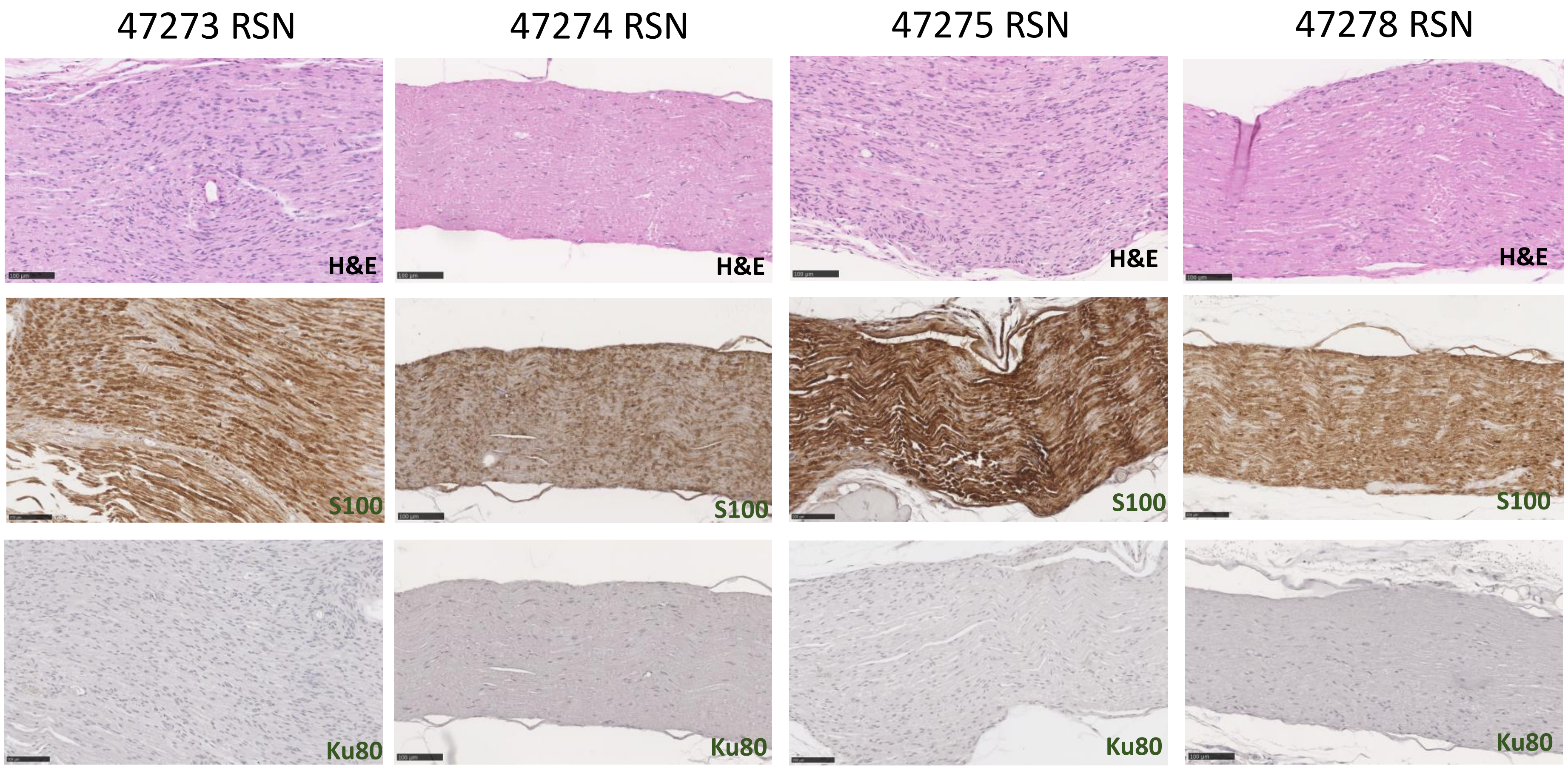
47273 RSN
47274 RSN
47275 RSN
47278 RSN
H&E
H&E
H&E
H&E
S100
S100
S100
S100
Ku80
Ku80
Ku80
Ku80

### Figure S1

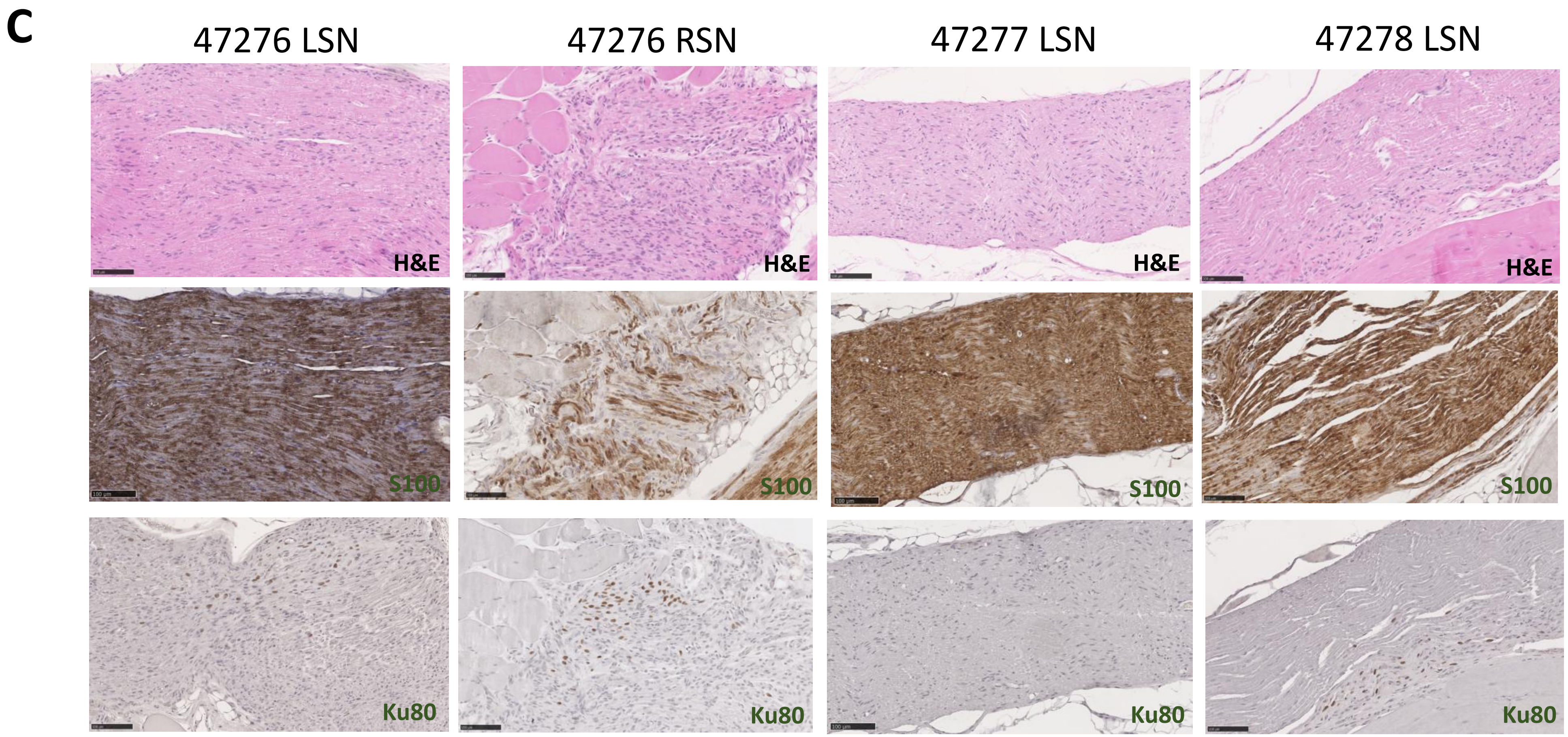

C
47276 LSN
47276 RSN
47277 LSN
47278 LSN
H&E
H&E
H&E
H&E
S100
S100
S100
S100
Ku80
Ku80
Ku80
Ku80

### Figure S1

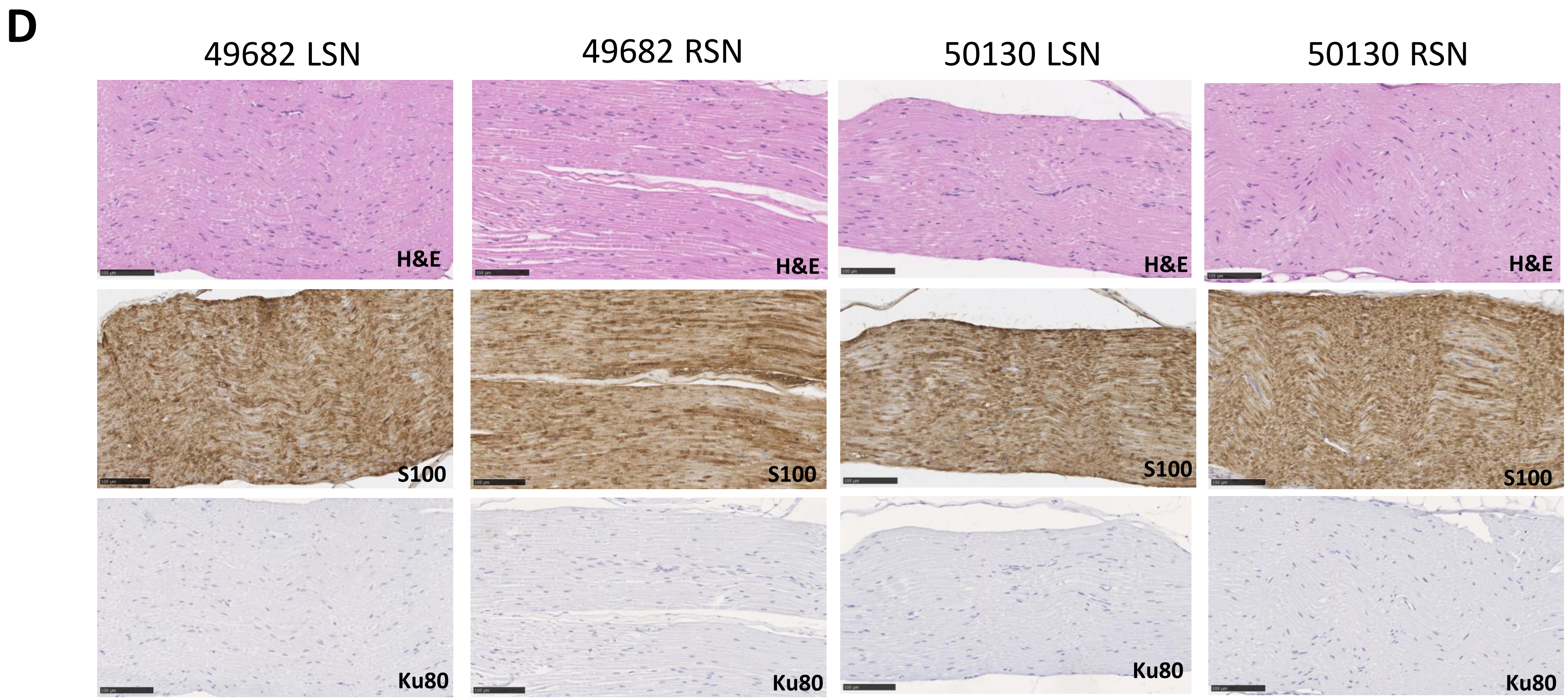
D
49682 LSN
49682 RSN
50130 LSN
50130 RSN
H&E
H&E
H&E
H&E
S100
S100
S100
S100
Ku80
Ku80
Ku80
Ku80

### Figure S1

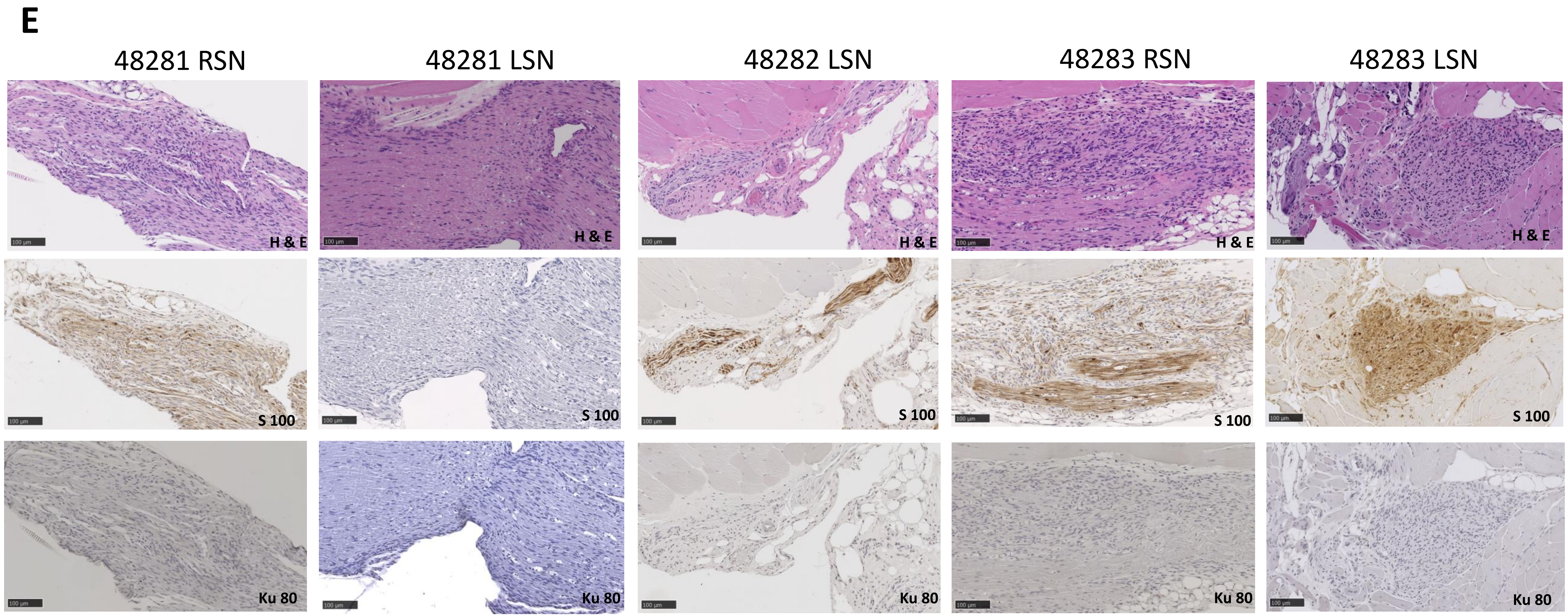
E
48281 RSN
48281 LSN
48282 LSN
48283 RSN
48283 LSN
H & E
S 100
Ku 80

### Figure S1

## F

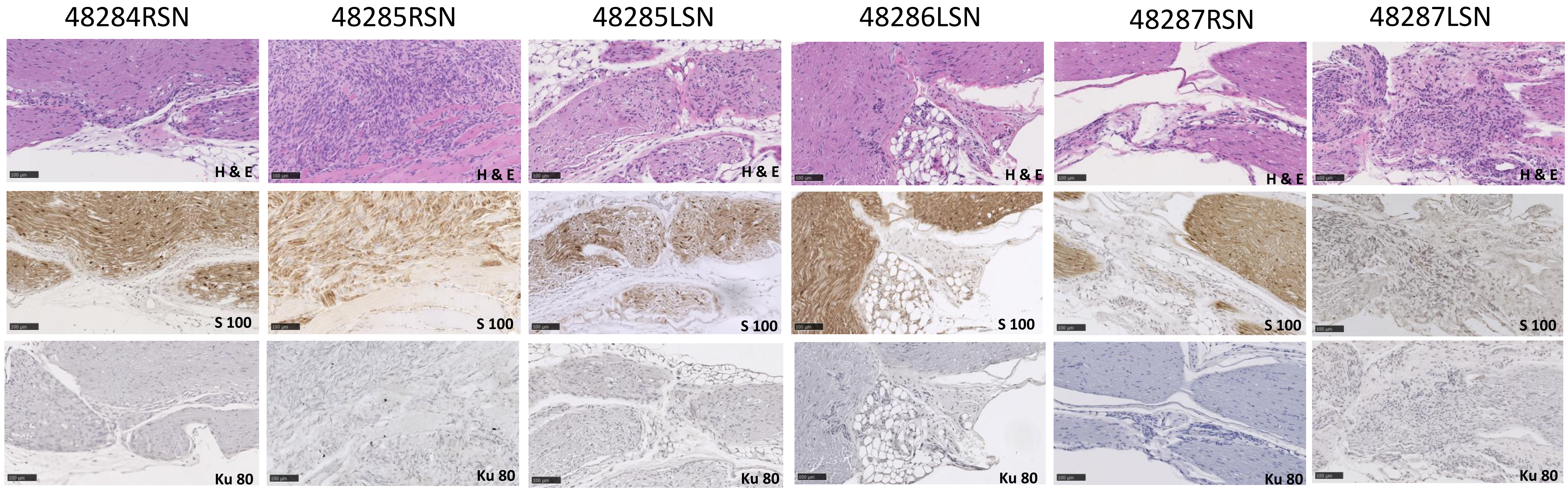

48284RSN
48285RSN
48285LSN
48286LSN
48287RSN
48287LSN
H & E
S 100
Ku 80

### Figure S2

## A

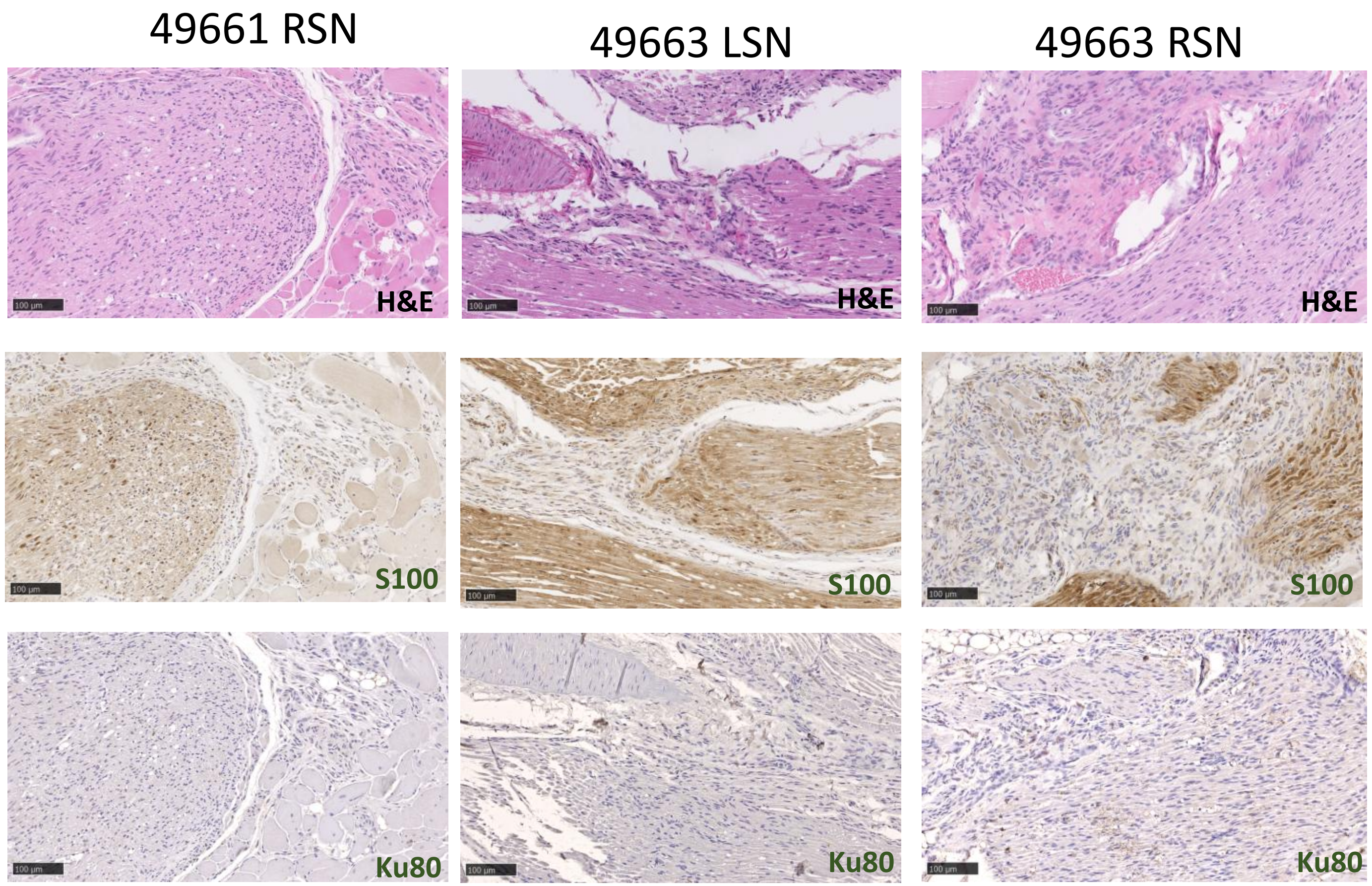

49661 RSN
49663 LSN
49663 RSN
H&E
H&E
H&E
S100
S100
S100
Ku80
Ku80
Ku80

### Figure S2

**B**

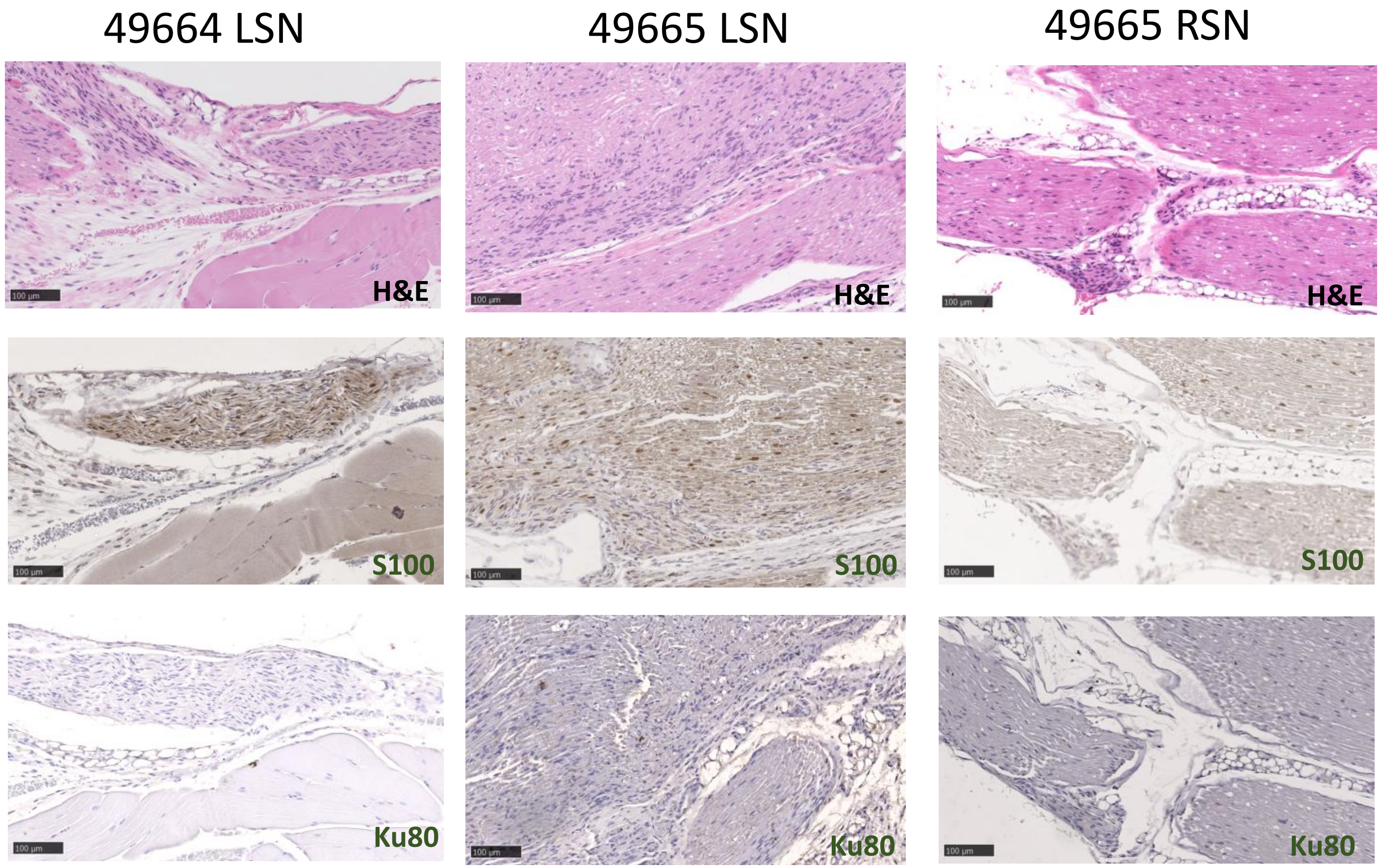
49664 LSN
49665 LSN
49665 RSN
H&E
H&E
H&E
S100
S100
S100
Ku80
Ku80
Ku80

### Figure S2

## C

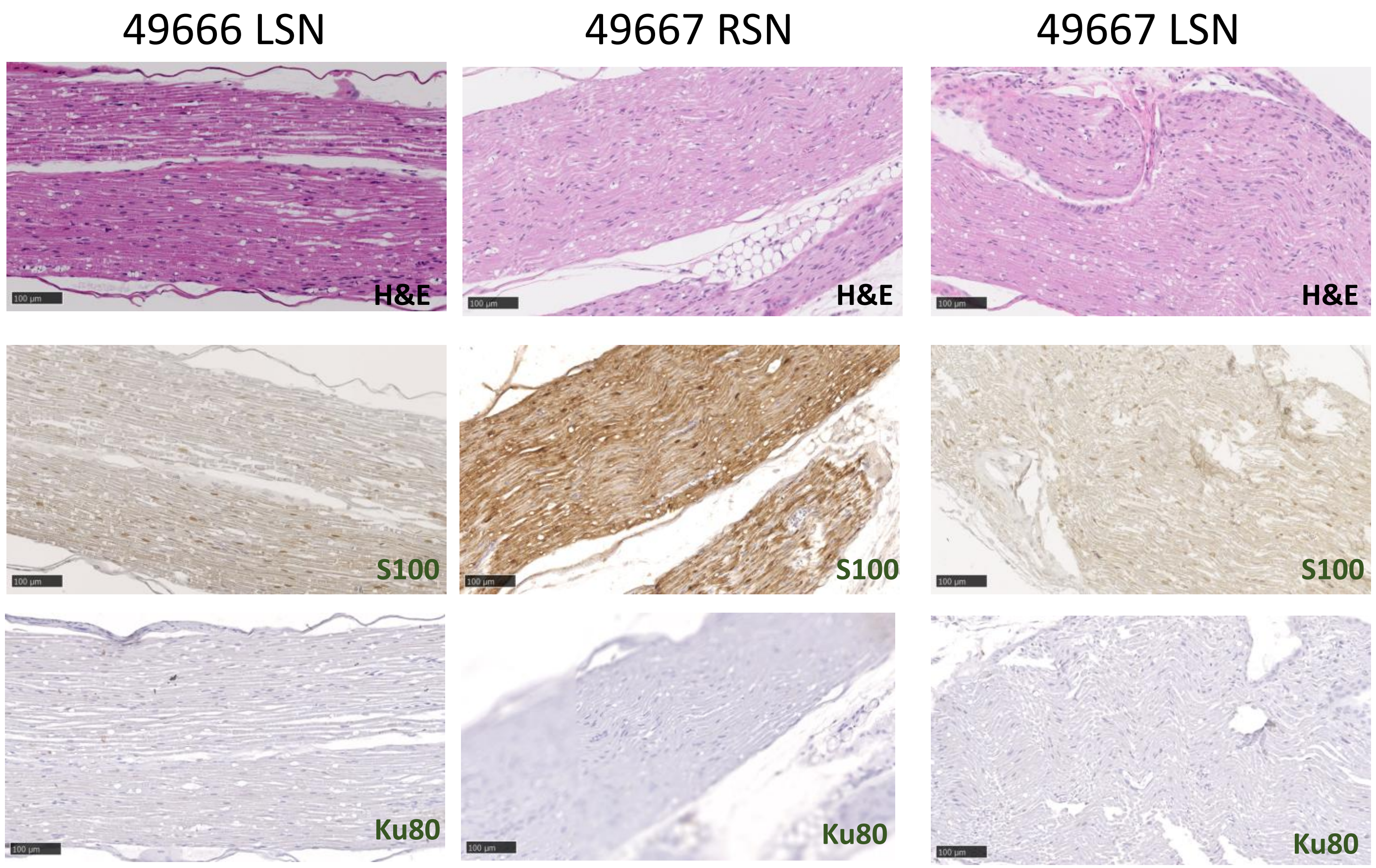
49666 LSN
49667 RSN
49667 LSN
H&E
H&E
H&E
S100
S100
S100
Ku80
Ku80
Ku80

### Figure S2

## D

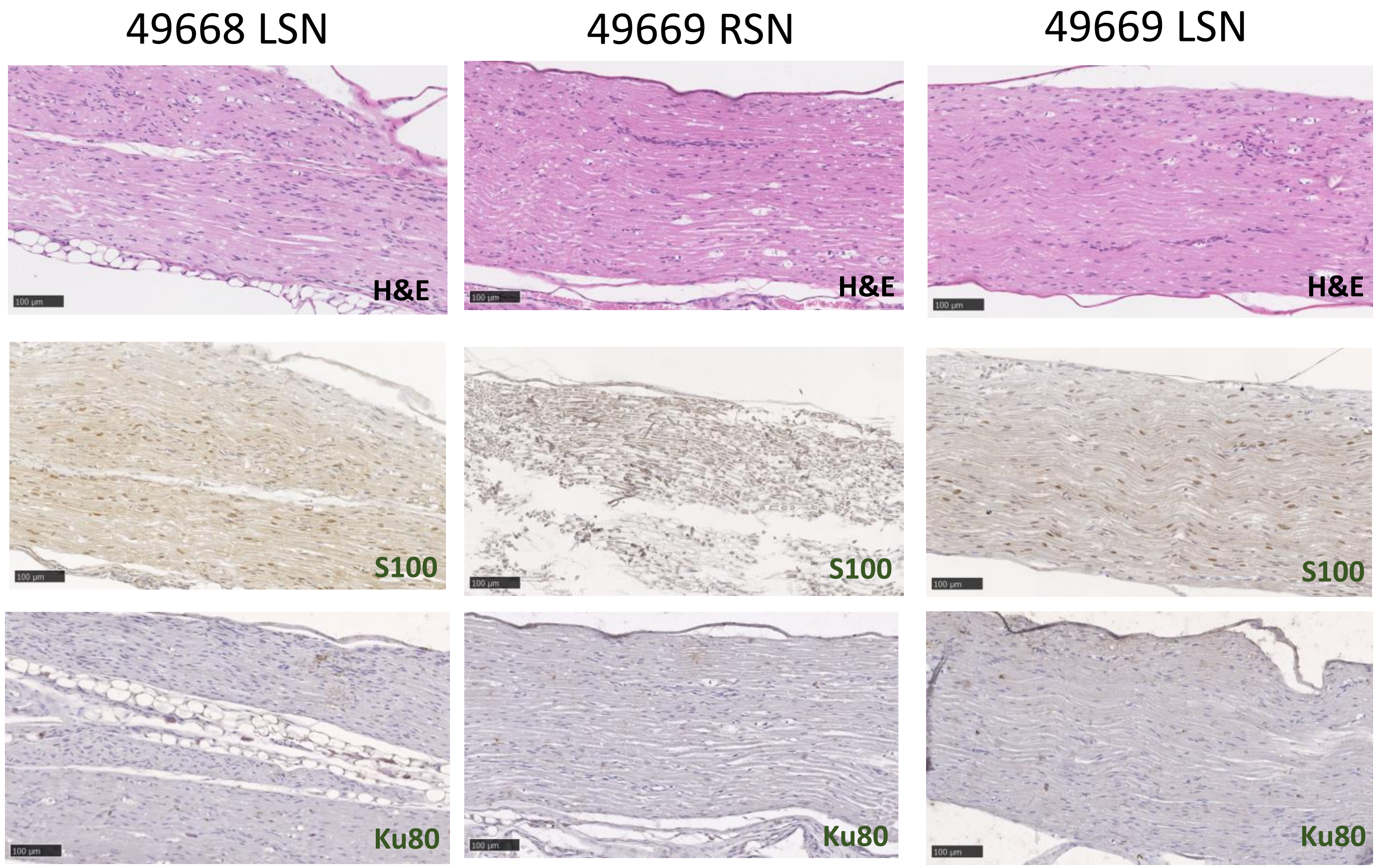

49668 LSN
49669 RSN
49669 LSN
H&E
H&E
H&E
S100
S100
S100
Ku80
Ku80
Ku80

### Figure S2

## E

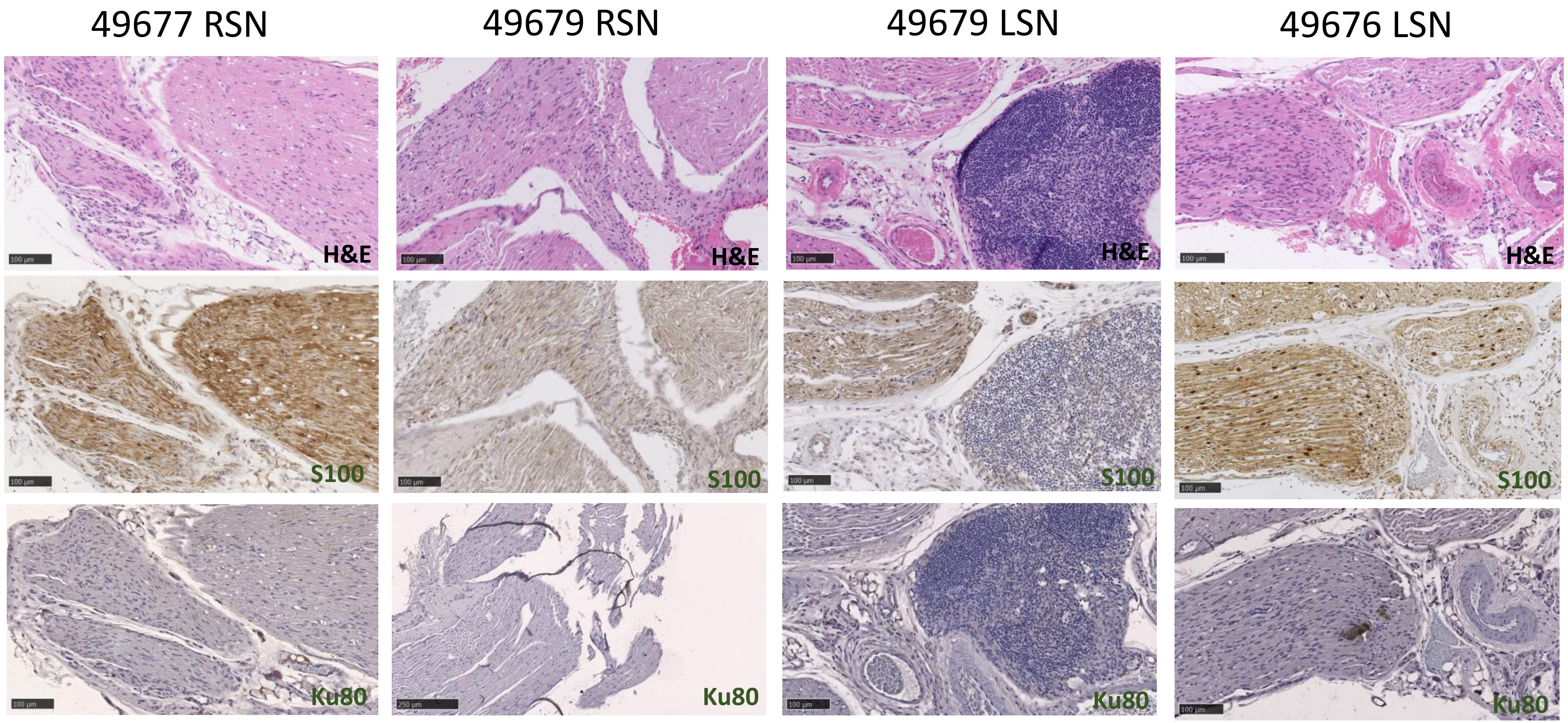
49677 RSN
49679 RSN
49679 LSN
49676 LSN
H&E
H&E
H&E
H&E
S100
S100
S100
S100
Ku80
Ku80
Ku80
Ku80

### Figure S3

## A

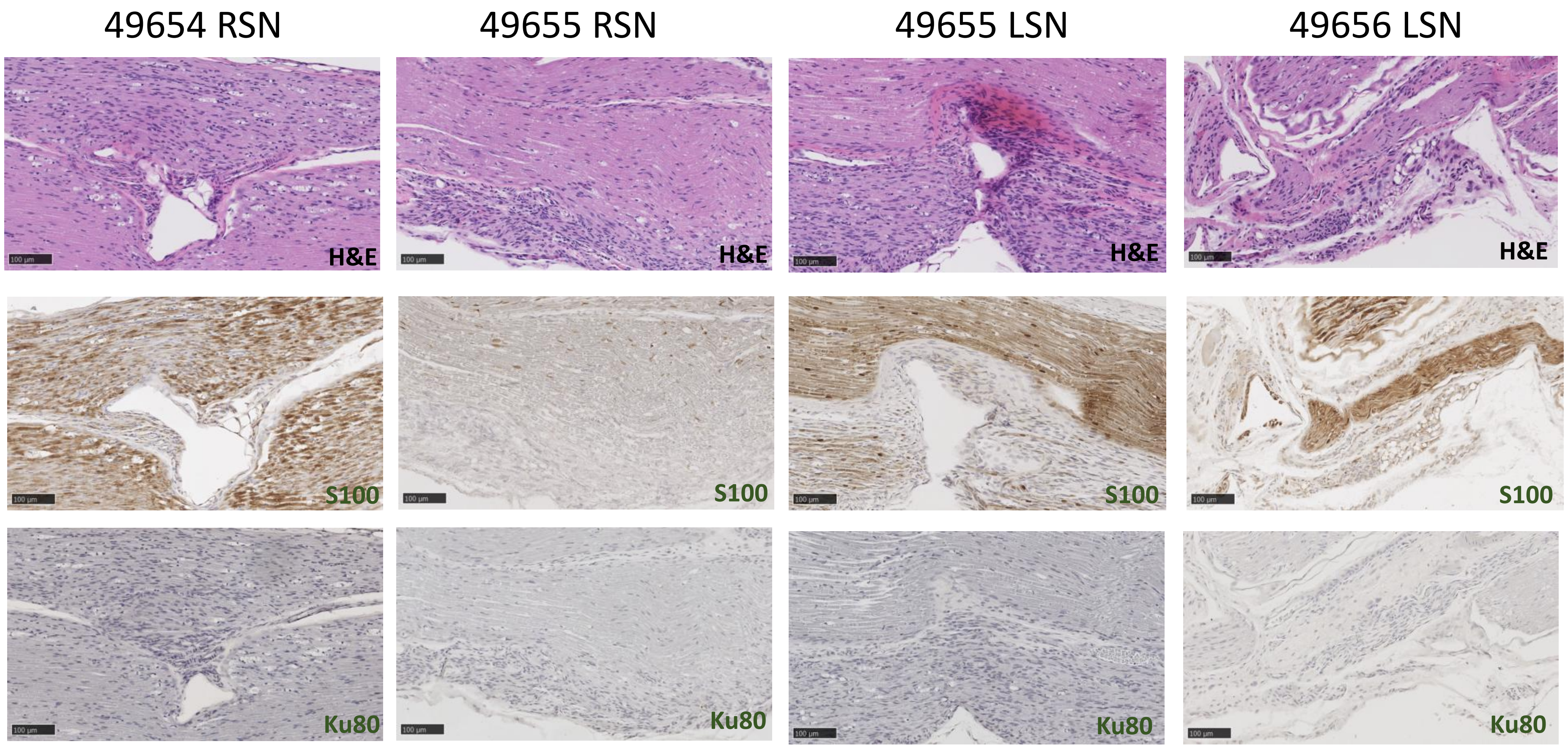

49654 RSN
49655 RSN
49655 LSN
49656 LSN
H&E
H&E
H&E
H&E
S100
S100
S100
S100
Ku80
Ku80
Ku80
Ku80

### Figure S3

## B

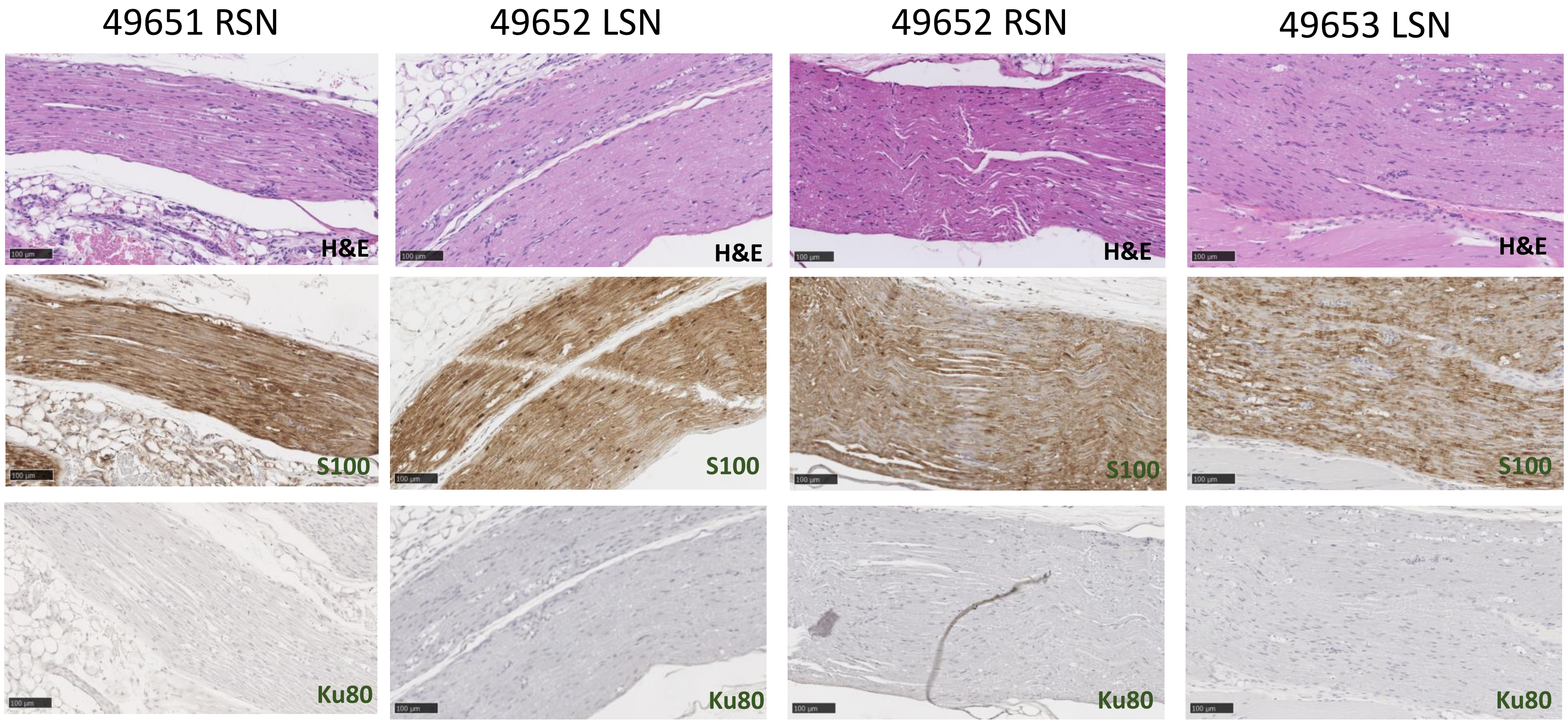

49651 RSN
49652 LSN
49652 RSN
49653 LSN
H&E
H&E
H&E
H&E
S100
S100
S100
S100
Ku80
Ku80
Ku80
Ku80
